# Genomes of the extinct Sicilian wolf reveal a complex history of isolation and admixture with ancient dogs

**DOI:** 10.1101/2022.01.21.477289

**Authors:** Marta Maria Ciucani, Jazmín Ramos-Madrigal, Germán Hernández-Alonso, Alberto Carmagnini, Sabhrina Gita Aninta, Camilla Hjorth Scharff-Olsen, Liam Thomas Lanigan, Ilaria Fracasso, Cecilie G. Clausen, Jouni Aspi, Ilpo Kojola, Laima Baltrūnaitė, Linas Balčiauskas, Jane Moore, Mikael Åkesson, Urmas Saarma, Maris Hindrikson, Pavel Hulva, Barbora Černá Bolfíková, Carsten Nowak, Raquel Godinho, Steve Smith, Ladislav Paule, Sabina Nowak, Robert W. Mysłajek, Sabrina Lo Brutto, Paolo Ciucci, Luigi Boitani, Cristiano Vernesi, Hans K. Stenøien, Oliver Smith, Laurent Frantz, Lorenzo Rossi, Francesco Maria Angelici, Elisabetta Cilli, Mikkel-Holger S. Sinding, M. Thomas P. Gilbert, Shyam Gopalakrishnan

## Abstract

The Sicilian wolf represented the only population of wolves living on a Mediterranean island until the first half of the twentieth century (1930s-1960s) ^1–7^. Previous studies hypothesised that they remained isolated from mainland wolves from the end of the Last Glacial Maximum (LGM) ^8,9^, until human persecutions led them to extinction ^1–7^.

There are only seven known Sicilian wolf specimens from the 19th and 20th century preserved in museums in Italy and recent morphometric analyses assigned them to the new subspecies *Canis lupus cristaldii* ^10^. To better understand the origins of the Sicilian wolf, and its relationship to other wolf populations, we sequenced four whole genomes (3.8×-11.6×) and five mitogenomes. We investigated the relationship between Sicilian wolves and other modern breeds to identify potential admixture. Furthermore, considering that the last land-bridge between Sicily and Italy disappeared after the LGM ^11^, around 17 kya, we explored the possibility that the Sicilian wolf retained ancestry from ancient wolf and dog lineages. Additionally, we explored whether the long-term isolation might have affected the genomic diversity, inbreeding levels and genetic load of the Sicilian wolf.

Our findings show that the Sicilian wolves shared most ancestry with the modern Italian wolf population but are better modelled as admixed with European dog breeds, and shared traces of Eneolithic and Bronze age European dogs. We also find signatures of severe inbreeding and low genomic diversity at population and individual levels due to long-term isolation and drift, suggesting also low effective population size.

## Results and Discussion

We sampled the seven available museum specimens of Sicilian wolf (Sicilian wolf information presented in Table S1), and successfully recovered four nuclear genomes and five mitogenomes. Poor DNA preservation hindered recovery of whole genome sequences in three samples and mitochondria of two. Raw reads were mapped to the dog reference genome (CanFam 3.1) ^12^, yielding average genomic depth of coverage ranging between 3.8× and 11.6× while the coverage on the mitochondrial DNA spanned between 19.7× and 1239.2× (Table S1). The analyses of the mapped reads and the damage pattern confirmed the authenticity of the historical DNA sequencing (Figure S1).

For comparative purposes we resequenced 33 modern wolf genomes (3.66×–41.9×) from across Europe. Further, we resequenced three Cirneco dell’Etna dogs (2×–2.6×), an old Sicilian hunting breed, that could have been in contact with Sicilian wolves. All modern samples were also mapped to the dog reference genome. To avoid biases introduced by ancient DNA damage, we limited all variant based analyses to transversion polymorphisms.

We determined the genetic sex of four Sicilian wolves, by comparing the average genomic coverage with the coverage of the X chromosome. The samples Sic1, Sic2 and Sic3 were identified as males (XY) while Sic7 was shown to be female (XX) (Table S1).

### Population structure and admixture of the Sicilian wolf

We first investigated the placement of the Sicilian wolves among the global dog and wolf diversity by computing a multidimensional scaling (MDS) analysis. We included previously described genomic groups of present-day wolves (Eurasian and North American), present-day and ancient dogs, and five ancient wolves that lived in Siberia during the Pleistocene ^13–15^. The first dimension separated dogs from wolves while the second dimension split European wolves from the rest of the Eurasian, North American and Pleistocene wolves. The results reveal a pattern of genetic isolation and drift of the Sicilian wolves analysed in this study (Figure 1A) which are placed in the upper right quadrant of the plot, within the European wolf diversity, in proximity to the Italian wolves, but shifted towards the cluster of dogs (Figure 1A and B). When restricting the MDS analysis only on wolves, the Sicilian wolf is represented even more differentiated and drifted compared to the Italian wolves and the rest of the European diversity (Figure S2A).

**Figure 1:**
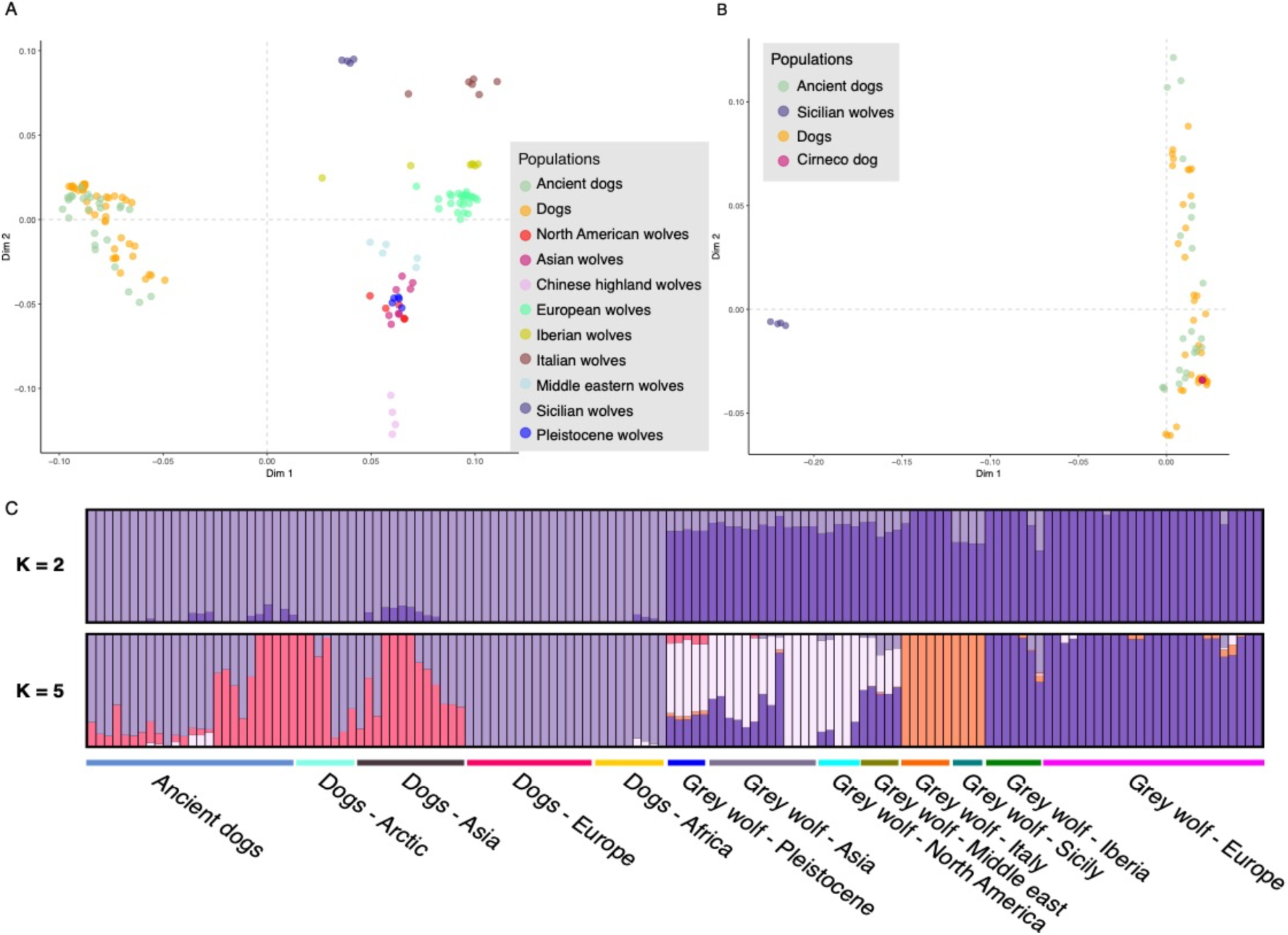
Population structure of the Sicilian wolves retrieved from genomic data. A) Multidimensional scaling plot presenting the Sicilian wolves in the ancient and modern wolves and dogs diversity. B) Multidimensional scaling plot with the Sicilian wolves among the diversity of modern and ancient dogs. C) Admixture plots showing two and five ancestral components (Ks).

We then performed an admixture analysis on the genomic data to determine the ancestry of the Sicilian wolf population using ADMIXTURE ^16^ (Figure 1C). When considering only the first two ancestral components (K=2) the Sicilian wolves are modelled as a mixture of wolf and dog components. Increasing the ancestral components up to five (K=5), the Sicilian wolves are represented by the same ancestral component as the Italian wolves. However, when considering seven ancestral components (K=7), there was further separation of the Sicilian wolves which were assigned to a different and unique component (blue) that remains consistent up to K=10 (Figure S2 B). These results confirmed that the genomic differentiation of the Sicilian wolves, supported also by the MDS plots, is consistent with the long-term geographic isolation and the extreme genetic drift that this population suffered in the past.

We investigated maternal lineage of the Sicilian wolves generating a mitochondrial phylogeny using RAxML-ng ^17^ including all the wolves and dogs in our dataset and using the coyote as the outgroup. The Maximum-Likelihood mitochondrial phylogeny is consistent with previous publications ^8,9^ and four out of the five Sicilian wolves (Sic1, Sic2, Sic3 and Sic7) cluster together in the phylogeny as a sister clade to wolves from Romania and Slovakia (Figure S2 D). They also fall within the same clade as wolves from Italy, Bulgaria, and Poland. The last sample, Sic4, falls within the dog mitochondrial diversity, as shown in a previous work based on the d-loop region ^8^. Unfortunately, we did not recover sufficient nuclear data from Sic4 to further investigate its ancestry. However, the sample was labelled as *Canis lupus*, but morphological analysis identified dog features, suggesting a hybrid origin. Our mitochondrial phylogenetic analysis placing Sic4 within dogs, is in agreement with the hybrid origin or, alternatively, it might suggest a potential mislabelling.

### The dual wolf-dog ancestry in the Sicilian wolves

Previous analyses based on the mitochondrial DNA haplogroups ^8,9^ clustered the Sicilian wolves together with both Northern European/Siberian Pleistocene and South-eastern European modern wolves. To test these previous results on the nuclear genomes and identify which were closest populations to the Sicilian wolves we performed an *Outgroup f3* analysis using Admixtools ^18^. Our results show that Northern European and Siberian Pleistocene wolves, together with East Eurasian, American and Middle Eastern wolves, share the least genetic drift with Sicilian wolves (Figure 2A and Figure S3A-D), indicating that they are the most distant populations. Instead, our results show that the Sicilian wolf is more closely related to modern Italian wolves, ancient European and modern Italian dogs (Figure 2A). In contrast, when comparing these results to the f3-statistics performed on Italian and Iberian wolves (Figure S3 E-F) we notice that the Sicilian wolf shares a similar or even higher level of drift than what was estimated for other individuals of the Italian and Iberian populations. These observations could be explained by the Sicilian wolf being highly differentiated due to long-term isolation.

**Figure 2:**
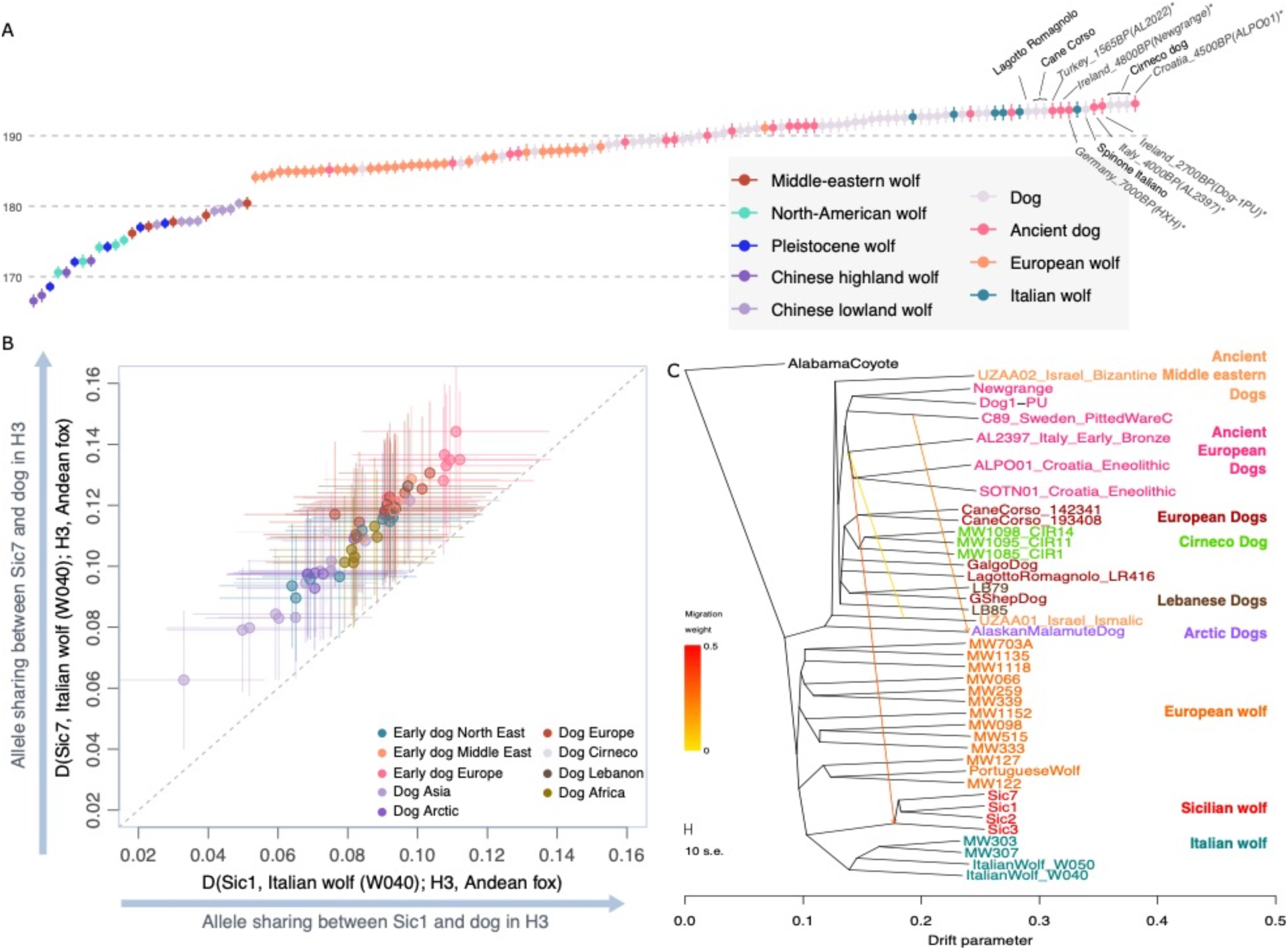
Relationship between Sicilian wolf and other populations/lineages (retrieved from genomic data). A) Outgroup *f3-statistics* showing the genetic affinity of the Sicilian wolf (Sic1) with other dogs and wolves. The plot shows the Siberian Pleistocene, North American, Middle Eastern and Asian wolves to be the most distant population from the Sicilian wolves. European wolves show slightly higher *f3* values. Italian wolves, European modern and ancient dogs show the highest *f3* values identifying these populations as the closest to the Sicilian wolves. B) *D-statistic* in the form of D(Sic1, Italian wolf, *Dogs*, Andean fox) - x axis - and D(Sic7, Italian wolf, *Dogs*, Andean fox) - y-axes - computed using qpDstats of Admixtools. All tests resulted in significant Z scores (| Z | ≥ 3.3). C) Treemix output showing 3 migration edges. The first high weight arrow shows migration from southern European ancient dogs into the Sicilian wolf, a second migration edge goes into the Alaskan Malamute from a Swedish (C89) Mesolithic dog. The third migration edge brings a Middle eastern dog component into southern European ancient dogs. TreeMix residuals of the graph with three migration edges are shown in Figure S5.

Together with the peculiar placement of the Sicilian wolves in the nuclear phylogeny (Figure S2 C) and admixture analysis, the *f3* results suggest the Sicilian wolf shares ancestry with both dogs and wolves.

Outgroup *f3*, computed between every pair of Sicilian wolves and dogs indicate an affinity between Italian dogs such as Spinone, Cane Corso and Cirneco with Sicilian wolves (Figure 2A), suggesting shared ancestry. Of all the modern dog breeds included in the analyses, the Cirneco dog is the closest modern dog to all Sicilian wolves (Figure S3 A-D). We hypothesise that these results could be driven by recent dog introgression due to the numerous stray dogs present on the island.

Interestingly, we also found stronger genetic affinity between the Sicilian wolves and ancient Eneolithic and Bronze Age dogs from Europe (∼5000–3000 BP - eg. Croatian ALPO01, Italian AL2397, Irish Dog-1PU and Newgrange, see Table S2, Figure 2A) than with modern European dogs. In particular, the Sicilian wolf sample Sic1 shares the most drift with ancient European dogs followed by Italian modern dogs. A similar pattern was observed in the other Sicilian wolves (Sic2, Sic3 and Sic7) (Figure S3 A-D). This genetic affinity with ancient European dogs could be due to: 1) introgression with ancient dogs, or 2) the Sicilian wolf being closer to a now extinct ancient wolf lineage that contributed to ancient European dogs. This last hypothesis could be supported by other studies, based on mitochondrial DNA ^19,20^, suggesting a genetic continuity with two Late Pleistocene wolves from Italy and dogs (ancient and modern). Therefore, to test these hypotheses, we computed *D-statistics* using qpDstats from Admixtool ^18^ to assess allele sharing between the Sicilian wolves, modern and ancient dogs, and wolves. Specifically, we computed all combinations of *D-statistics* of the form D(Sicilian wolf, Italian wolf; *Dogs*, Andean fox). We find the Sicilian wolf shares significantly more alleles with all dogs compared to the modern Italian wolf (|*Z*|≤3.33). Furthermore, our results show that the ancient European dogs yield the largest values of D (Figure 2B and Figure S4A-B).

The outgroup *f3* and *D-statistics* results led us to focus our analyses on the investigation of the relationship between ancient dogs and Sicilian wolves. To disentangle the historical relationships between Sicilian wolves, modern and ancient dogs, we used TreeMix ^21^ to build a graph incorporating different numbers of admixture events. The TreeMix graph without migration edges is consistent with the nuclear phylogeny (Figure S2 C) placing the Sicilian wolves as basal to all modern and ancient dogs (Figure S5 A). Once migrations were incorporated into the analyses, the resulting graphs placed the Sicilian wolves as sister clade to the Italian wolves while drawing a high weight (weight = 0.322216) ancestry contribution from Croatian Eneolithic dogs going into the base of their clade (Figure 2C and S5).

Furthermore, since in the *D-statistics* tests D(*European ancient or modern dogs, Dogs/Wolves*; Sicilian wolf, Andean fox), the Sicilian wolf (Sic1) resulted to be equally distant to Italian wolves, ancient and modern European dogs (Figure 3A-B) we proceeded to model the ancestry proportions of the Sicilian wolf, taking into account gene flow from other populations. To do so we used qpGraph ^18^ (Figure 3C and Figure S4C) in which ancient and modern dogs, Sicilian, Italian and Iberian wolves were included. Consistent with results from the *f3, D-statistics* and TreeMix, this analysis indicates that the Sicilian wolf ancestry can be modelled as a mixture of ancient dogs (31%) and Italian wolves (69%).

**Figure 3:**
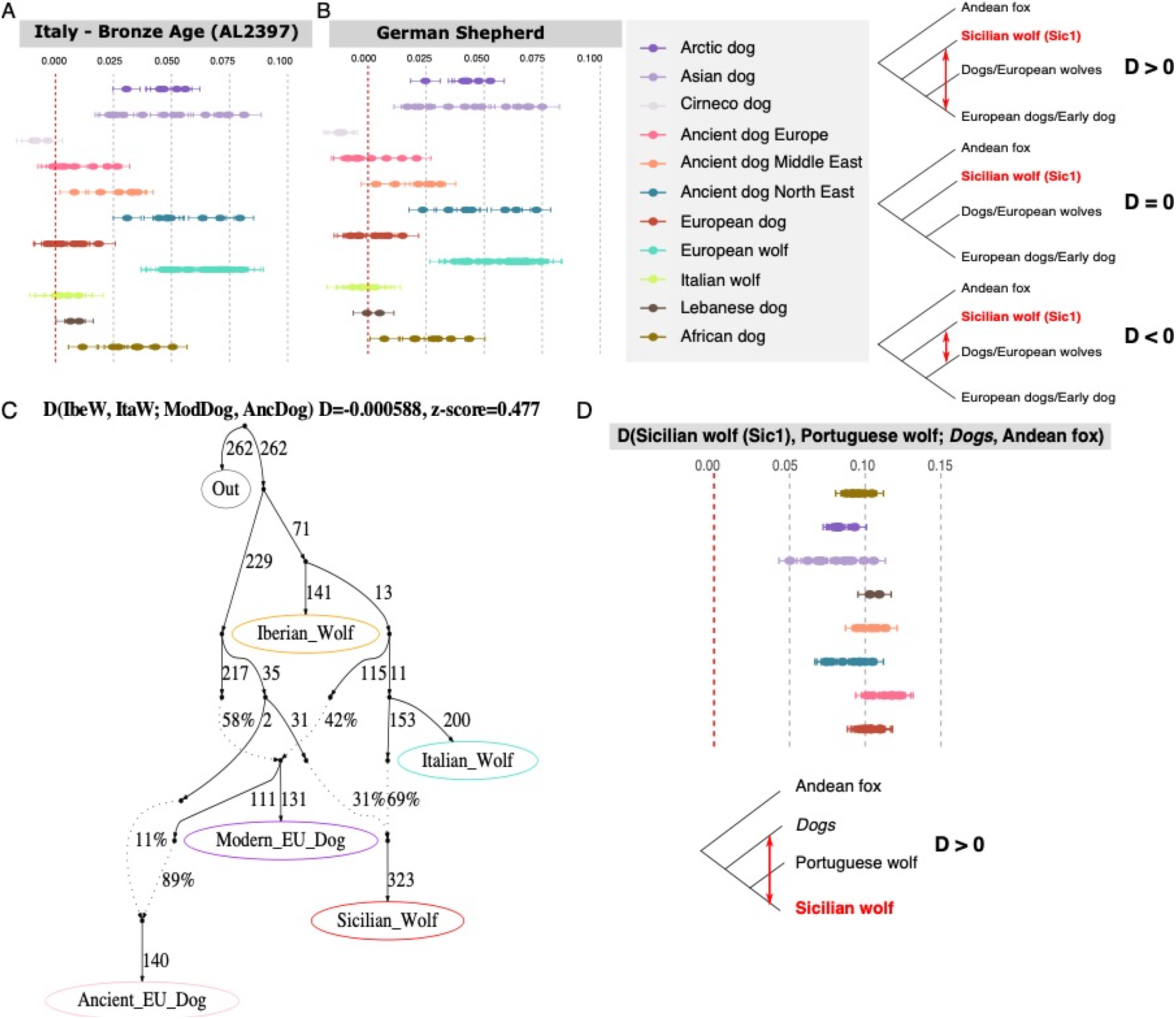
Relationship between Sicilian wolf and dogs. A-B) *D-statistics* computed using qpDstats of Admixtools. A) *D-statistic* in the form of D(Ancient dog (Italy AL2397), *Wolves/Dogs*; Sic1, Andean fox). B) *D-statistic* in the form of D(German Shepherd, *Wolves/Dogs*; Sic1, Andean fox). C) qpGraph admixture graph modelling the Sicilian wolf as a mixture between a common ancestor with the Italian wolf (69%) and ancient European dogs (31%). D) *D-statistics* investigating the directionality of gene flow between dogs and Sicilian wolves. The assumption behind this test is that if all the *D-statistics* of the D(Sicilian wolf, Portuguese wolf, *Dogs*, Andean fox) are significant with a Z-value < 3 the result is BABA meaning gene flow from dogs into Sicilian wolf. On the contrary if some tests were significant but negative resulting in ABBA that could be an indicator that the gene flow could have been from Sicilian wolf into certain dogs leading us to investigate the next D test.

Lastly, we investigated the directionality of gene flow between Sicilian wolves and dogs by estimating the *D-statistics* in the form D(Sicilian wolf, Portuguese wolf, *Dogs*, Andean fox). We expected all results to be positive if the Sicilian wolf carries dog ancestry. Conversely, if the gene flow goes from Sicilian wolf into specific dog breeds, we would expect only those to yield significant values. Our result consistently shows positive results for all the Sicilian wolves suggesting, therefore, gene flow from dogs into Sicilian wolves (Figure 3D).

Overall, the *D-statistics* and *f3-statistics* results show all four Sicilian wolves to have a similar ancestry pattern, where they carry a substantial dog ancestry component. A previous study on Sierra Morena wolves (*C. l. signatus*) found that they also carried a high dog ancestry proportion, up to a third of their genome ^22^. This population shared similar characteristics as the Sicilian wolf, being a small, isolated and declining population in a human-modified landscape. Our results further suggest that these features could contribute to increased hybridization due to the combined effects of stray dogs and dwindling population sizes of wolves. This is not surprising considering that the samples used in this study represent the last individuals of Sicilian wolves and it is likely that the degree of hybridisation with modern dogs was significantly lower in the past. We can see that a high degree of hybridisation with modern dogs is also present in one of the last specimens of Japanese wolves (*C. l. hodophilax*) which became extinct at the same time as the Sicilian wolf ^23,24^.

### The effects of island isolation on the Sicilian wolves

We then explored the insular effect and the consequences of the population decline in the last few hundred years on the genomic diversity of the Sicilian wolf population.

Compared to other populations, we find that the Sicilian wolves have a low population level nucleotide diversity (π) and at the individual level this is confirmed by very low heterozygosity (Figure 4 A-B). These results are expected in populations that are particularly affected by founder effect, long-term isolation and small effective population sizes ^25–28^.

**Figure 4:**
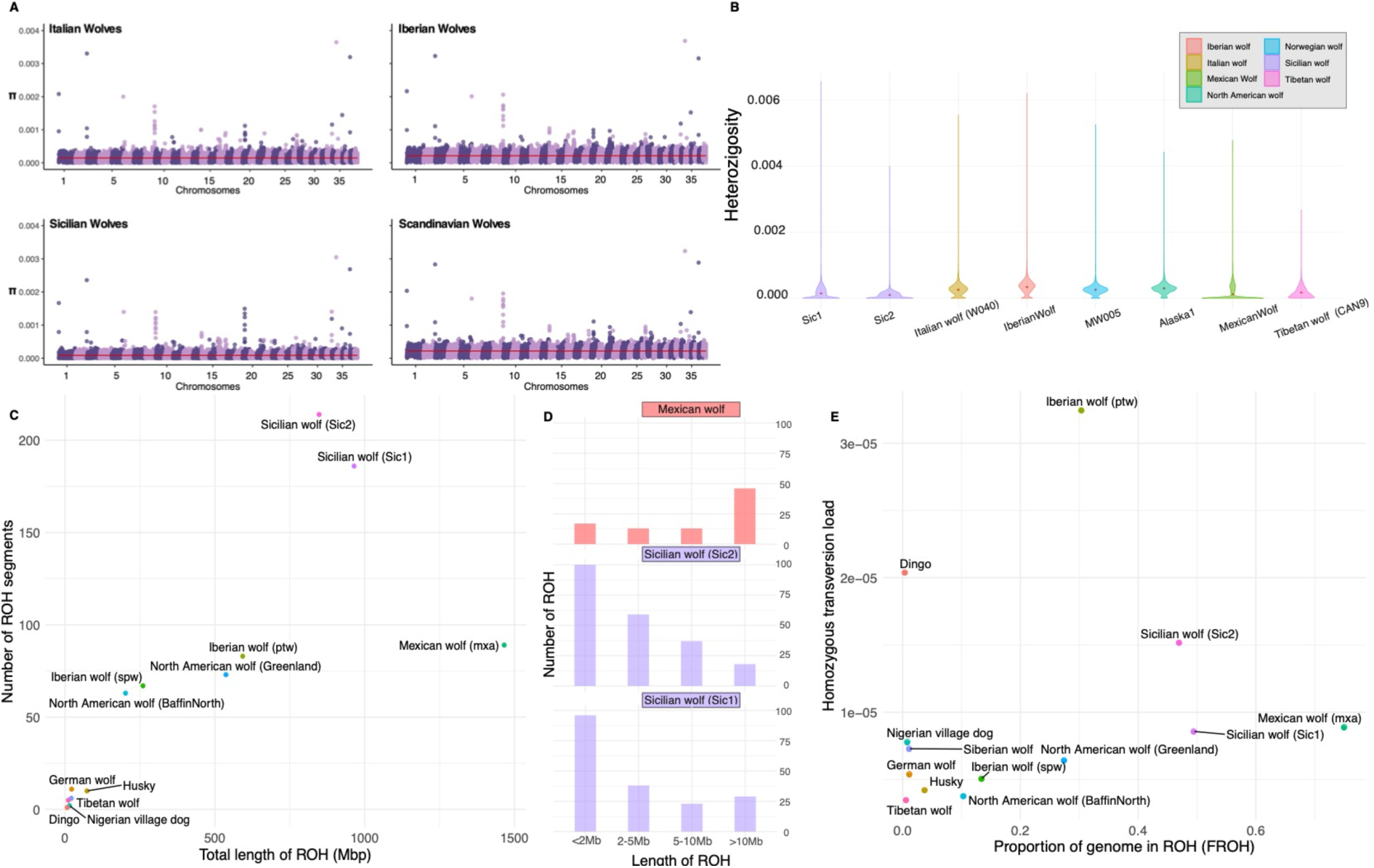
Island isolation effect. A) Population level nucleotide diversity of Sicilian, Italian, Iberian and Scandinavian wolves. The chromosomes and the nucleotide diversity values are shown on the x and y axis respectively. The red line represents the mean diversity value. B) Heterozygosity in sliding windows (without including repetitive regions) for different wolf samples show reduced levels of heterozygosity in the Sicilian wolves Sic1 and Sic2. C) Individual runs of homozygosity represented as the number of ROH segments on the y-axis and total length of ROH (Mbp) on the x-axis. D) Histogram of the number of small, medium and long ROH segments. The Sicilian wolves (Sic1 and Sic2) show remarkably high numbers of ROH, particularly short ones. E) Genetic load represented as the number of transversion sites that are homozygous is shown on the y-axis compared to the proportion of the genome in ROH (FROH). The Sicilian wolves seem to have a higher transversion load compared to other wolf populations.

Furthermore, we find that the Sicilian wolves have a mean length of runs of homozygosity (ROHs) of ∼ 5Mb, with a high proportion of their genome in ROHs, *ca* 40% and 35% for Sic1 and Sic2 respectively. While the proportion of the genome with long runs of homozygosity (ROH ≥ 2Mb) is still particularly elevated (*ca* 32%), the number of short and medium size ROH segments is higher than the ones observed in other highly and recently inbred population such as the Mexican wolf (*C. l. baileyi*) (Figure 4D). The Sicilian wolves also contain a relatively higher mutation load with respect to their inbreeding extent (Figure 4E). Overall, these results indicate that the Sicilian wolf population already at the end of the 19th century was suffering from the effect of small population size leading to mating between closely related individuals. However, short ROH segments could indicate that the inbreeding happened in the past, rather than towards the time of extinction, as a result of perhaps previous persecutions.

To conclude, our study showed that the Sicilian wolf population was strongly affected by long-term insular isolation with particularly low population and individual levels of diversity and a high proportion of the genome in ROH.

The Sicilian wolves also shared common ancestry with modern Italian wolves and dogs. The similarity between Italian and Sicilian wolves, and the evidence for long-term isolation (i.e. long-term inbreeding), raised the question about the timing of colonisation of Sicily by the ancestor of the Sicilian wolf. In the past, due to the climatic changes during the Pleistocene, Sicily was occasionally connected to Italy through a land bridge on the Strait of Messina. This patch of land disappeared, *ca* 17 kya, at the end of the last glacial period, due to increased sea levels, and since then the Strait has always been characterised by strong currents prohibiting the crossing in the past ^11^. Therefore, even if wolves have proven to be able to swim ^29^ it is unlikely that a substantial number of Italian wolves colonised Sicily by sea after the end of the LGM.

Our results, showing the Sicilian wolf closest to the Italian wolves, support that around the time of Sicilian colonisation, the founder population present in the Italian Peninsula was already differentiated from the other European wolf populations (including the Iberian and Balkan wolves)^30^. Additionally, previous studies, based on the whole nuclear genome, suggested that contemporary wolf populations shared a common ancestor between 32 kya and 13 kya^13,14,31^. We suggest that the last common ancestor between the Italian and the Sicilian wolves colonised Sicily through the land bridge between 25 kya and 17 kya^9^.

Furthermore, from our analyses we detected an excess of allele sharing between Sicilian wolves and European dogs, particularly with ancient Eneolithic and Bronze dogs. This indicates that Sicilian wolves sequenced could be the result of admixture, likely limited in the island, with ancient European dogs in the past, and potentially with additional modern dog admixture up to the time of extinction.

This study shows that the Sicilian wolf population survived until the 20th century preserving traces of the ancient dog lineage that was widespread in Europe until 3-4000 years ago ^32^. Among all other extant and extinct European wolf populations that have been genetically characterised until now, the Sicilian wolf is unique in showing such a strong signal of genetic affinity with ancient European dogs.

## Acknowledgements

This work was supported by ERC Consolidator Award 681396 Extinction Genomics, DNRF143 Center for Evolutionary Hologenomics, and the Norwegian Environment Agency (project 18088069). We would like to thank Luca Sineo for helping collect the Sicilian wolf samples and Domenico Tricomi, together with the owners of the Cirneco dell’Etna dogs for their collaboration in this project. We are grateful to the curators of the museums who allowed us to sample the historical Sicilian wolves: Paolo Agnelli of the Museum of Natural History of the University of Florence, zoology section “La Specola”, Fabio Lo Bono of the Civic Museum “Baldassare Romano” in Termini Imerese (PA), Enrico Bellia of the Museum of Zoology “P. Doderlein”, SIMUA, Palermo (PA) and Ferdinando Maurici and Fabio Lo Valvo from Regional Museum of Terrasini (PA). We also thank the staff of the Danish National High-throughput Sequencing Centre and BGI Denmark for their support in data generation and the Danish National Supercomputer for Life Sciences at the DTU National Lifescience Center at Technical University of Denmark (DTU) for facilitating the data analysis process. The National Science Centre, Poland supported S. Nowak (grant No. 2020/39/B/NZ9/01829) and R.W. Mysłajek (grant No. 2019/35/O/NZ8/01550). G.H-A. is supported by the Consejo Nacional de Ciencia y Tecnología from Mexico (CONACyT 576734). R.Godinho is supported by the Portuguese Foundation for science and Technology (DL57/2016/CP1440).

## METHOD DETAILS

### Description of the historical Sicilian samples

The sample Sic1 (cat number C11875) was a petrous bone from a wolf skull preserved at the Museum of Natural History, Section of Zoology ‘La Specola’, University of Florence, Florence, Italy. The sample still has the original tag from the 19th century, and it belongs to a wolf killed in 1883 at Vicari (PA). The sample Sic2 (cat number AN/855) is represented by a petrous bone from a wolf skull belonging to a juvenile individual from 1879 preserved at the Museum of Zoology ‘Pietro Doderlein’ of the University of Palermo, Palermo, Italy. The sample Sic3 (cat number M/18) is a mounted specimen of an adult male. This individual was killed in Sicily, presumably around 1870-1880 as documented by an old picture in the museum records. The specimen is preserved in the Museum of Zoology ‘Pietro Doderlein’ of the University of Palermo, Palermo, Italy. The individual Sic4 (cat number 9) is represented by a hide labelled as an adult wolf that was shot in 1924 at Bellalampo (PA), and it is preserved in the Regional Museum of Terrasini (Palermo), Italy. The last sample, Sic7 (cat number NA) is a hide dated around 1880-1920 and is preserved at the Museum of Termini Imerese (Palermo), Italy.

### Data generation for the historical Sicilian samples

The Sicilian wolf samples were processed under strict clean laboratory conditions at the GLOBE Institute, University of Copenhagen. DNA extractions were performed by following the silica-based protocol described in Dabney et al (2013) ^33^ for the bone samples and Campos & Gilbert ^34^ in case of keratin samples (hides and claws). Both bone and keratin samples were digested using 1 mL digestion buffer. The extracts were purified using modified PB buffer (Qiagen), washed twice with PE buffer (Qiagen) and eluted twice in 20 μL of buffer EB - with 10 minutes of incubation time at 37°C. The concentration of each extract was checked on a Qubit Fluorometer (ThermoFisher Scientific) in ng/μL and on the Tapestation (Agilent Technologies) for concentration and fragment size.

BGI libraries for the samples Sic1 and Sic4 were constructed following Carøe et al. ^35^ and Mak et al. ^36,37^ while BEST Illumina libraries for Sic2, Sic3 and Sic7 were built following Carøe et al. using Illumina adapters ^35^. Libraries were prepared using up to 32 μL of DNA in a final reaction volume of 80 μL.

The appropriate number of cycles for the amplification were determined using Mx3005 qPCR (Agilent Technologies) in which 1 μL of SYBRgreen fluorescent dye (Invitrogen, Carlsbad, CA, USA) was loaded in 20 μL indexing reaction volume using also 1 μL of template, 0.2 mM dNTPs (Invitrogen), U/μL AmpliTaq Gold DNA polymerase (Applied Biosystems, Foster City, CA, USA), 2.5 mM MgCl2 (Applied Biosystems), 1X GeneAmp® 10X PCR Buffer II (Applied Biosystems), 0.2 μM forward and reverse primers mixture ^36^, and 16.68 μL AccuGene molecular biology water (Lonza, Basel, CH). qPCR cycling conditions were 95°C for 10 minutes, followed by 40 cycles of 95°C for 30 seconds, 60°C for 30 seconds, and 72°C for 45 seconds.

The libraries were amplified using PfuTurbo Cx HotStart DNA Polymerase (Agilent Technologies) and Phusion® High-Fidelity PCR Master Mix with HF buffer (New England Biolabs Inc). The amplification was performed in 50 μL PCR reactions that contained:

- 14 μL of purified library, 0.1 μM of each forward (BGI 2.0) and custom made reverse BGI primers, 2x Phusion® High-Fidelity PCR Master Mix with HF buffer and 8.6 μL AccuGene molecular biology water.
- 10 μL of purified library, forward and reverse primers, 2.5U/L PfuTurbo Cx HotStart DNA Polymerase, 0.4 BSA, 1 μL of dNTPs (25 μM), 1 μL of Buffer 10X and 30.6 μL AccuGene molecular biology water.

PCR cycling conditions for libraries amplified using Phusion® High-Fidelity PCR Master Mix with HF buffer were: initial denaturation at 98°C for 45 seconds followed by 18 to 20 cycles of 98°C for 20 seconds, 60°C for 30 seconds, and 72°C for 20 seconds, and a final elongation step at 72°C for 5 minutes. Amplified libraries were then purified using 1.5x ratio of SPRI beads to remove adapter dimers and fragments smaller than 90-100bp and eluted in 50 μL of EB (Qiagen) buffer after 10 minutes incubation at 37°C. The samples Sic1 and Sic4 were sequenced using BGIseq platform while the samples Sic2, Sic3 and Sic7 were pooled together and sequenced on Novaseq6000 Illumina platform, S2 flow-cell PE50.

### Data generation for modern samples

#### Cirneco dell’Etna buccal swab samples - DNA Sample Collection, Storage and Extraction

SK-1S Isohelix buccal swabs for non-invasive DNA collection were used to sample three Cirneco dell’Etna individuals (MW1085_CIR1, MW1095_CIR11, MW1098_CIR14). The dogs were sampled by their owners at least 30 minutes after eating and to avoid contamination gloves and face masks were used. The swabs were inserted into the dogs’ mouth, rubbed on their cheek for *ca* 1 minute and placed back into the collector tube together with the SGC-50 Isohelix Dri-capsule to preserve and stabilize the DNA at room temperature during shipping. Once received in the Modern DNA labs of the GLOBE Institute (University of Copenhagen) the samples were stored at -20°C.

Buccal swab samples were placed in 2 mL Eppendorf tubes and were extracted using a modified version of the DNeasy® Blood and Tissue Kit (Qiagen). In each tube were added 380 μL of ATL Buffer and 20 μL of Proteinase K (Roche). The samples were placed in a thermomixer for 1 hour at 56°C. The lysate was transferred in a different tube and 400 μL of AL Buffer and 400 μL of 96% Ethanol were added. The extraction reaction was then spun down in a DNeasy Mini column and the filter was washed with 500 μL of Buffer AW1 and 500 μL of Buffer AW2. The DNA was eluted twice using 50 μL of AE Buffer directly onto the DNeasy Mini column membrane.

#### Modern Italian wolves tissue samples - DNA Sample Collection, Storage and Extraction

The Italian wolf samples were collected around 2010 and 2012 from road-killed individuals populating the Simbruini Mountain Range Regional Park and National Park of Abruzzo in the Central Apennines (See Table S2). The muscle samples were stored in ethanol at -20°C and subsequently processed in the “Modern DNA laboratory” at the Fondazione Edmund Mach (FEM). Small pieces of tissue of around 25 mg were extracted using the DNeasy Blood and Tissue Kit (Qiagen) with an overnight digestion at 56°C. The elution was conducted at the GLOBE Institute (University of Copenhagen) using two washes of 50 µL of AE buffer, with 10 minutes of incubation at 37°C. Until the elution, samples were stored at -20°C inside the DNeasy Mini spin columns.

#### Modern wolves - DNA Sample Collection, Storage and Extraction

Modern wolf tissue and blood samples from several locations in Europe (Table S2) were extracted using Thermo Scientific KingFisher instrument and following the manufacturer’s protocol. The samples were then checked for concentration with a Qubit Fluorometer (ng/μL) All the extracts - with the exception of the modern Italian wolves and the Cirneco dell’Etna - were sent to BGI Copenhagen for library build and sequenced on ⅛ of a lane each on DNBSEQ PE150.

#### Library build, amplification and sequencing of modern Italian wolves and Cirneco dogs

Extracts were fragmented in the Covaris LE220 plus Focused-ultrasonicator with the parameters set for getting 350-bp fragment length. The extracts were diluted to obtain 100 ng concentration and BGI libraries for the Italian wolves and Cirneco dogs were constructed following Carøe et al. ^35^ and Mak et al. ^36^ using 10 µM adaptors. Libraries were purified using MinElute columns using PE buffer (Qiagen) and eluted in 60 µL of EB buffer.

The PCR mixture for the Italian wolf libraries consisted on: 20 μL of purified library, 0.2 μM of forward and reverse BGI primers, 2.5 U/μL PfuTurbo Cx HotStart DNA Polymerase, 10 μL of Buffer 10X, 0.08 mg/mL BSA, 0.5 mM of dNTPs (25 μM) and 61.2 μL AccuGene molecular biology water (Lonza, Basel, CH). The Cirneco dog libraries were amplified using 20 μL of purified library, 0.2 μM of forward and reverse BGI primers, 0.05 U/μL of AmpliTaq Gold (Thermo Fisher Scientific, USA), 0.4 mg/mL BSA, 0.2 mM of dNTPs (25 μM), 10 μL of Buffer 10X, 2.5mM of MgCl2 and 50.2 μL AccuGene molecular biology water in a total volume of 100 μL reaction.

The amplification of the Italian wolves was performed as follows: initial denaturation at 95°C for 2 minutes followed by 10 to 12 cycles of 95°C for 30 seconds, 60°C for 30 seconds, and 72°C for 110 seconds, and a final elongation step at 72°C for 10 minutes. PCR cycling conditions for Cirneco dog libraries were: initial denaturation at 95°C for 10 minutes followed by 12 cycles of 95°C for 20 seconds, 60°C for 30 seconds, and 72°C for 45 seconds, and a final elongation step at 72°C for 5 minutes. The Italian wolves and Cirneco dogs were sequenced on ⅛ of a lane each on MGIseq2000 PE150 and DNBSEQ PE150 respectively.

### Data processing and historical DNA authentication

Short sequencing reads from modern, historical, and ancient samples were mapped to the dog reference genome (CanFam3.1)^12^ using Paleomix v.1.2.13 pipeline ^38^. First, adaptor sequences were removed using AdapterRemoval 2.0 ^39^ with default settings and BWA v0.7.12 *backtrack* ^40^ was used to perform the alignment of reads to the dog genome (bwa seed was disabled) setting the minimum base mapping quality to 0 to retain all the reads in this step. In further computational steps reads with mapping quality lower than 20 or 30 will be discarded. Picard MarkDuplicates v2.9.1 ^41^ was used to filter out PCR duplicates and GATK v4.1.0 ^42,43^ was used to perform the indel-realignment step.

For the historical Sicilian samples in this study the post-mortem DNA profiles and misincorporation patterns were performed through mapDamage2.0 ^44^.

### Dataset

The dataset designed for this study is represented by 154 samples (Table S2), including one Andean fox (*Lycalopex culpaeus*)^45^, three golden jackals (*Canis aureus*) ^13,14,46^, four coyotes (*Canis latrans*) ^46,47^, four African golden wolf (*Canis lupaster*) ^46,48,49^, one Ethiopian wolf (*Canis simensis*) ^46^, 62 modern wolves (*Canis lupus*) from Eurasia and North America ^13,14,22,46,50–53^, 45 modern dogs ^12,14,45,51–59^ (see Table S2 for more information), 24 ancient dogs (*Canis lupus familiaris*) ^32,51,60–63^, five Pleistocene wolves ^15,51^ and five newly sequenced Sicilian wolves (*Canis lupus cristaldii*). In this dataset 33 modern wolves and three Cirneco dogs were newly sequenced as part of this study. See Table S2 for information regarding sample ID, location, coverage, project number and publication for each sample used.

### Variant calling

For each modern sample at each genomic site, we sampled a random read using ANGSD v0.931 ^64^ (-doHaploCall 1) from the reads with a minimum mapping quality of 30 (-minMapQ 30) and bases with minimum quality of 20 (-minQ 20). The following parameters were used in the command line: - minMinor 1 -maxMis 10 -skipTriallelic 1 -doMajorMinor 2 -C 50 -baq 1 -remove_bads 1 - only_proper_pairs 1. The output file was converted in Plink files (tped and tfam) using haploToPlink tool in ANGSD and used to create a list of variable sites to call in ancient and historical samples and outgroups. Also, in this case ANGSD with -doHaploCall option was used to call the haplotypes for the list of sites (option -s) that were variable across the modern samples. Transitions were discarded (- rmTrans 1) to reduce the aDNA derived error in the historical samples included in our dataset. The modern, ancient, and historical samples were merged and converted in Plink files. The resulting dataset consisted of 74.5 million SNPs that we were able to merge.

Plink v1.9 ^65^ was used to prune the panel from minimum allele frequency below 5% (-maf 0.05), missing data (N) and filter out all the variants with missing call rates below 25% (-geno 0.25). Variants in linkage disequilibrium (LD) within 10kb window size were also removed (--indep-pairwise 10kb 2 0.5). The remaining panel contained 2.3 million SNPs and it was used (or a subset of it) for the following analyses and named “haploid SNPs panel”.

### MDS

The SNPs panel created with ANGSD in the previous section that consist of 153 samples and a total of 2.3 million transversion sites was used to generate a multidimensional scaling plot (MDS) by estimating first the IBS (pairwise distance) matrix between samples using Plink v1.9 ^65^. The MDS plots were generated including all the wolf and dog samples, one including all the samples, one with only wolves and another one with only dogs and the Sicilian wolves.

### Admixture

The haploid SNPs panel including wolves and dogs was used as input in the ADMIXTURE ^16^ analysis to estimate the ancestry components on a subset of samples in the dataset. The outgroups were not included in this analysis. ADMIXTURE was run on 2 to 10 ancestry components (K2 to K10), and, for each K, 50 independent replicates were performed. The replicate with the best likelihood value was chosen for each K. Pong ^66^ was used to visualise the admixture plots.

### Mitochondrial and nuclear phylogenies

We built a mitochondrial phylogeny using all the samples in the dataset (n = 131) using the coyote as the outgroup. To generate the fasta files we extracted the chrMT from the bam file using Samtools ^67^, and built a majority consensus sequence using ANGSD (-dofasta 2). The mitochondrial sequences for each sample were combined and aligned using MUSCLE ^68^, and a ML tree was built with RAxML-ng ^17^ using the evolutionary model GTR+G and 500 bootstrap replicates. The resulting tree was visualised using FigTree v.1.4.4 ^69^.

To build the nuclear phylogeny on all the modern and ancient individuals in the dataset, we first used ANGSD through which we generated a consensus sequence for each genome using the dog reference genome (CanFam3.1) ^12^ as reference and sampling each base randomly (-dofasta1). Bases with base quality lower than 20 and reads with mapping quality lower than 20 were discarded (-minQ 20 -minmapq 20). The minimum coverage for each individual was set to 3x (-setminDepthInd 3), and the following additional filters were used: -doCounts 1 -remove_bads 1 -uniqueOnly 1 -baq 1 -C 50. We then selected only from autosomal chromosomes 1000 random regions, each 5000bp long, from the dog reference genome using BEDTools^70^ *random* with the following parameters: -l 5000 -n 1000. We then used Samtools^67^ to select from each sample the regions of interest and combined them into a multi-sequence alignment (MSA) fasta file. Each MSA files were used in IQ-TREE ^71^ v.2.1.2 to reconstruct the phylogeny using 1000 bootstrap replicates (-B 1000) with UFBoot2 ^72^, 1000 bootstrap replicates for SH-aLRT (-alrt 1000) and *ModelFinder Plus* ^73^ to identify the best evolutionary model for each region. The resulting 1000 trees were then concatenated in a single file and used as input in Astral-III ^74^ to estimate the nuclear phylogeny. FigTree ^69^ v.1.4.4 tool was used to visualise the species tree estimated with Astral-III.

### Outgroup three population test (Outgroup *f3-statistics*)

We calculated *outgroup f3* using qp3pop tool in Admixtools ^18^ v.5.1 to assess the shared genetic drift between the reference population A and B since the separation from the outgroup Andean fox (*A, B; AndeanFox*). A negative *f3-statistics* value with the correspondence Z-value less than -3 indicates that the population A is close to the population B. The higher the value the closer the 2 populations are to one another.

### Four-population test (*D-statistics*)

We computed *D-statistics* on the haploid dataset using qpDstats in Admixtools ^18^ v.5.1 to study how modern and ancient wolves and dogs are related to the Sicilian wolves.

The following forms of *D-statistics* were tested:

1. D(Sicilian wolf, Italian wolf; *Wolf/Dog*, Andean fox) (Figure 2B and Figure S4A);
2. D(ancient European breed, *Wolf/Dog*; Sic1, Andean fox) (Figure 3A);
3. D(modern European dogs, *Wolf/Dog*; Sic1, Andean fox) (Figure 3B);
4. D(*Dogs*, Cirneco dog; Sicilian wolf, Andean fox) (Figure S4B);
5. D(*Dogs*, Croatian Eneolitic dog (ALP01); Sicilian wolf, Andean fox) (Figure S4B);
6. D(Sicilian wolf, Portuguese wolf; *Dogs*, Andean fox) to determine which wolf populations have a closer relationship to dogs (Figure 3D). We assumed that if all d-stats are significant and positive the gene flow could be from dogs into Sicilian wolves. If some tests are not significant it could indicate that the gene flow is directed into certain dogs.

In all the D-statistic tests the outgroup (Andean fox) is placed in H4 therefore if the value of the *D-statistics* is different from 0 it means that there is allele sharing with one population from H1 or H2 to the population in H3. Specifically, if the value is negative with a Z-value smaller than -3 (|Z| < - 3) the result will be ABBA, implying allele sharing between populations H2 and H3. Instead, if the d value in positive – with a Z-value greater than 3 (|Z| > 3) – the results will be the opposite and the allele sharing will be between H1 and H3 (BABA).

### TreeMix

The software TreeMix ^21^ v1.13 was used to evaluate the admixture event from dogs into Sicilian wolves to understand, in combination with *f3* and *D-statistics*, the dog ancestral component into wolves. TreeMix was run on a subset of the haploid dataset including the four Sicilian wolves, Italian wolves, Iberian and other European wolves and ancient and modern dogs for a total of 37 samples. We restricted the analysis to sites without missing data resulting in a final dataset of 160,233 transversion sites. The program was run ten times (10 replicates) for each migration event (m) ranging from 0 to 4. The optional parameters -noss, -global and -k 500 were used and the tree was rooted using a coyote (Alabama coyote). The trees with the best likelihood for each migration event were selected using R ^75^ and plotted using the plotting_funcs.R provided by Treemix software. The topology of these clades remains overall constant from the 1st up to 4 migration events, and none of the inferred migrations involves the Cirneco dog and the Sicilian wolves.

### Admixture graphs

To model the relationships between the Sicilian wolf, modern wolves and dogs we estimated an admixture graph using qpGraph ^18^. We included samples from relevant groups: Italian wolf (MW303, MW307, ItalianWolf_W050, ItalianWolf_W040), Iberian wolf (PortugueseWolf, MW122, MW127), modern dog (BoxerDog, GalgoDog, GShepDog), ancient dog (ALPO01_Croatia_Eneolithic, SOTN01_Croatia_Eneolithic, AL2397_Italy_Early), Sicilian wolf (Sic1, Sic7, Sic2) and the Andean fox. First, we identified samples that could be grouped into each of the relevant groups using qpWave ^18^. By comparing two lists of populations (left and right populations) qpWave identifies the minimum number of migrations required from the right populations to define the ancestry of the left populations. As ‘left populations’ we tested all possible pairs of Italian wolves, Iberian wolves, modern European dogs, ancient dogs and Sicilian wolves. For the ‘right populations’ we used all other samples, except the ones included among the ‘left populations’ and using the Andean fox as fixed outgroup. qpWave was run using the ‘allsnps=YES’ option. Pairs of samples that were consistent with a single migration (p-value ≥ 0.05) were grouped into a population for the admixture graphs: Italian wolf (MW303, MW307 and ItalianWolf_W040), Iberian wolf (PortugueseWolf, MW122, MW127), ancient dog (ALPO01_Croatia_Eneolithic and AL2397_Italy_Early), modern European dog (GalgoDog and GShepDog) and Sicilian wolf (Sic1 and Sic7).

To efficiently explore the possible admixture graph models we used two approaches: qpBrute ^63,76^ and a ‘base graph’ approach as described in Ramos-Madrigal ^15^. For the qpBrute approach we created a configuration file for qpBrute and performed a heuristic search of the admixture graph space. For the ‘base graph’ approach we use *admixturegraph* R library ^77^ to list all possible graphs including the Italian wolf, Iberian wolf, ancient dog, and modern European dog, and fixed the Andean fox as an outgroup. We tested all possible graphs including 0, 1 and 2 migration edges. We then included the Sicilian wolf to the two graphs that fitted our data in terms of the Z-score of the worst fitting *F*-statistic and with a score that was not significantly different ^78^. The Sicilian wolf was included both as a non-admixed lineage and as an admixed lineage. We show the best graph from each approach from the graphs that fitted our data (Figures 3C and S4C).

### Nucleotide diversity and Heterozygosity in sliding windows

The genome-wide nucleotide diversity (π) was estimated for four wolf populations using 4 Sicilian wolves, 5 Italian wolves, 4 Scandinavian wolves and 6 Iberian wolves. Vcftools 0.1.16 ^79^ was run on the haploid SNPs panel with a window of 100kbp. Rstudio ^80,81^ and ggplot ^82^ were used to visualise the data for each population.

We used ANGSD to estimate the heterozygosity of each sample, by calculating the folded site frequency spectrum (SFS) on the autosomal chromosomes. A saf.idx file based on individual genotype likelihoods using GATK (-GL 2) was generated for each bam file (doSaf 1 -fold 1) and the transitions (rmtrans 1) and reads with quality score and bases with mapping quality lower than 20 (-minQ 20 - minmapq 20) were excluded. The dog reference genome was used both as reference and as ancestral (- ref and -anc options) and the repeat regions were masked using a repeat mask file (-sites). The chromosomes were partitioned into overlapping windows of size 1 Mb with a step size of 500 kb using the BEDTools ^70^ windows tool. Windows shorter than 1Mb at the end of the chromosomes were discarded. The SFS for each window was estimated using the realSFS utility tool provided in ANGSD and subsequently the final heterozygosity per window was calculated as the ratio of heterozygous sites/total sites. Rstudio, ggplot2 and dplyr ^83^ were used to visualise the heterozygosity level at each window in the form of a violin plot for each individual.

### ROHan

To assess the extent of recent inbreeding, we estimated the length and abundance of segments of runs of homozygosity (ROH) using ROHan on two genomes of Sicilian wolf (Sic1 and Sic2) with minimum 5x following the recommended coverage for ancient genomes ^84^. Before running ROHan, we created a bed file containing mappable regions in the reference dog genome (CanFam3.1) to account for divergence between the genomes of the wolf and the dog. Then, a file listing only the autosomal chromosomes was created to limit our analysis to the autosomal regions. Afterwards, to get the number of segregating sites expected in the ROH segments (--rohmu) required by ROHan, we estimated background heterozygosity from X chromosomes of male canids which found an average of 4 × 10^−5^ segregating sites. We also observed the deamination profile of the ancient genomes using bam2prof (included in ROHan) and determined that for most of the ancient bam files the damage levels were off at -q 20. The bam2prof results were then used on the ROHan call for the Sicilian wolf genomes (-- deam5p and --deam3p).

We calculated the number of segments (NROH), mean length (LROH), and total sum of runs of homozygosity (SROH) in the autosomal region of each sample from the .hmmrohl output of ROHan using an R script. The genomic coefficient of inbreeding was calculated in the same script by dividing SROH with the total number of validated sites by ROHan available on the .hEst file. The presented plot only includes the middle mid-point estimates from the three mid-point estimates provided by ROHan (min, mid, and max).

### Transversion load

To assess the abundance of deleterious mutations in the wolves genome, we used the conservation scores from SIFT, following the pipeline of Greer (2020) ^85^. More specifically, we obtained the conservation scores from VEP Ensembl, selected several taxa to obtain ancestral allele of positions, and used this along with the genotype probability values from each sample to calculate mutation load. We consider mutation load as the sum of conservation scores where there are homozygous transversion weighted by the genotype probability of that site divided by the sum of genotype probability of homozygous transversion and total the whole k positions in the genome:

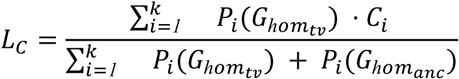

As the SIFT score for deleterious allele tend to approach 0 and the score is per allele, we calculated load by selecting two kinds of homozygous derived genotypes that are transversions from ancestral alleles, marked by 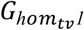 and 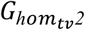 with their respective SIFT scores *S*_*1*_ and *S*_*2*_.

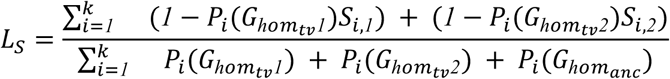

