## Supplemental File for "Genomes of the extinct Sicilian wolf reveal a complex history of isolation and admixture with ancient dogs"

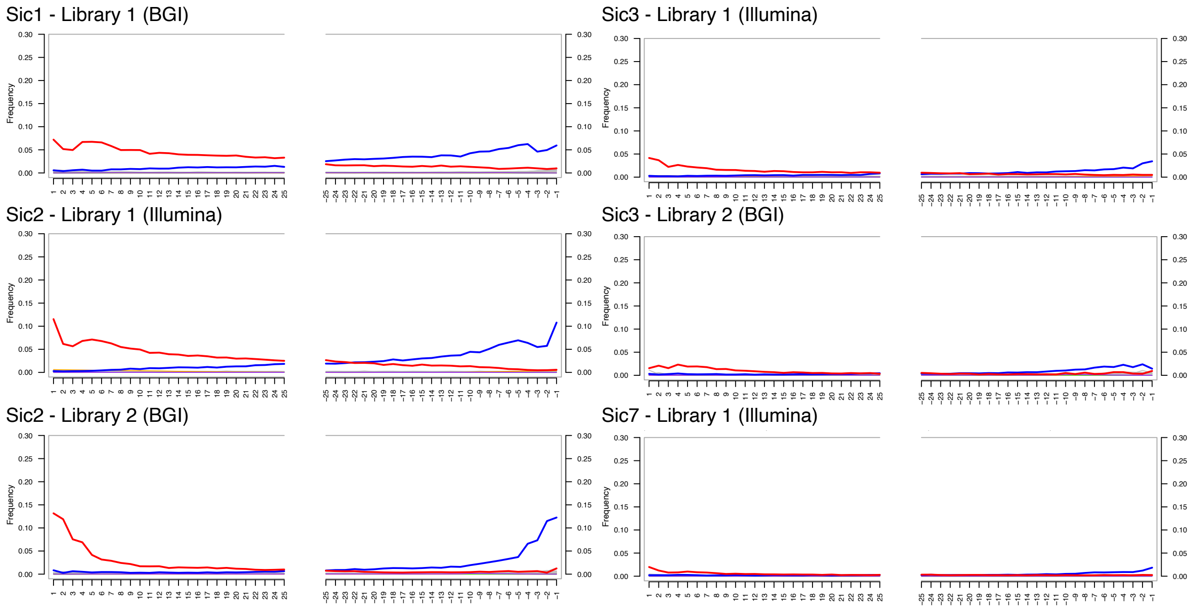


**Figure S1: Sicilian wolves' misincorporation plots from mapDamage [1].**

Nucleotide misincorporation patterns of the Sicilian wolf samples mapped to the dog reference genome (CanFam3.1)[2]. Two plots are shown for Sic2 and Sic3, one for the Illumina libraries and one for BGI libraries. For the samples Sic1 and Sic7 only the Illumina libraries plots are shown.

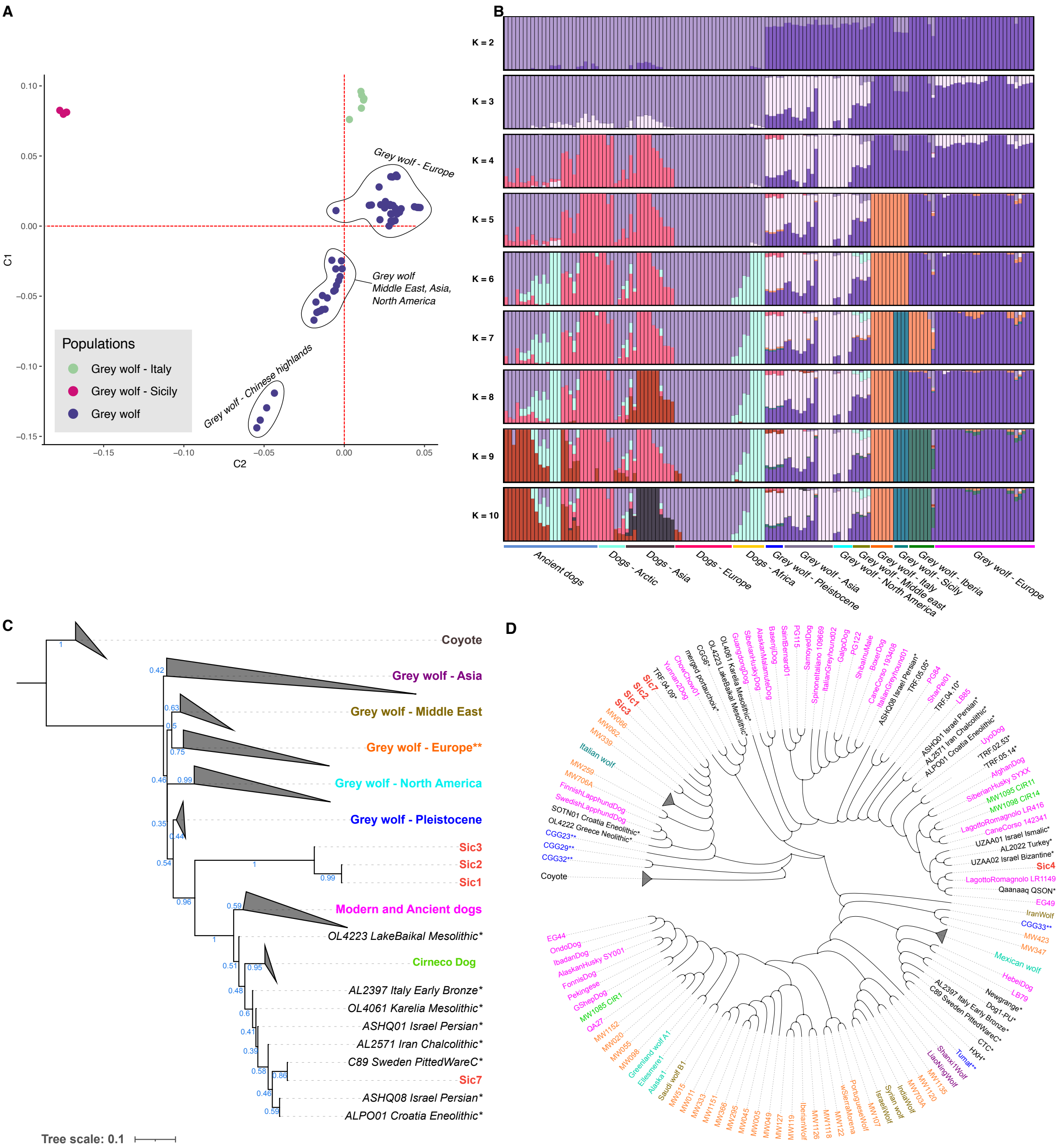


**Figure S2: Population structure and phylogenies.**

A) Multidimensional Scaling (MDS) plot of wolves. Grey wolves from Eurasia and North America are represented in purple while Italian and Sicilian wolves are represented in light green and pink respectively.

B) Admixture proportions for a range of 10 ancestral components (K=10) were estimated using ADMIXTURE [3]. For each component, 50 independent replicates were performed and the ones with the better likelihood were selected.
The cross validation values for each Ks are the following: K2= 0.86411; K3=0.86986; K4=0.89088; K5=0.91350; K6=0.94008; K7=0.96207; K8=0.99581; K9=1.02498; K10=1.06985. Since the lowest value represents the best fitting plot, K2 seems to fit our data best.

C) Nuclear phylogeny of wolves and dogs using coyotes as outgroup. The phylogeny was computed by ASTRAL-III [4] on 1000 trees generated with IQtree [5] v.2.1.2 using 1000 bootstrap replicates and *ModelFinder Plus* [6] to identify the best evolutionary model for each region. The Sicilian wolves are represented in bold red, while the star (*) represents ancient samples. Modern and ancient dogs are represented in pink, Pleistocene wolves in blue, North American wolves in turquoise, European wolves in orange, Middle Eastern wolves in sand and Asian wolves in purple.
(**) Note that the Italian wolves are collapsed within the European wolf clade.

To determine the phylogenomic placement of the Sicilian wolves among the wolf and dog's diversity we built a tree based on autosomal chromosomes using ASTRAL-III [4] by combining 1000 trees from randomly selected regions across the genome. The estimated species tree was rooted using the coyote as the outgroup. The tree shows that Sicilian wolf samples do not cluster within the modern European wolves clade, instead they are placed basal to all dogs included in the tree.

D) Mitochondrial phylogeny of wolves and dogs using the coyote as an outgroup. The Sicilian wolf is represented in red. The label of each sample is colored based on its group or geographic location according to the same color pattern shown in Figure S2 C.

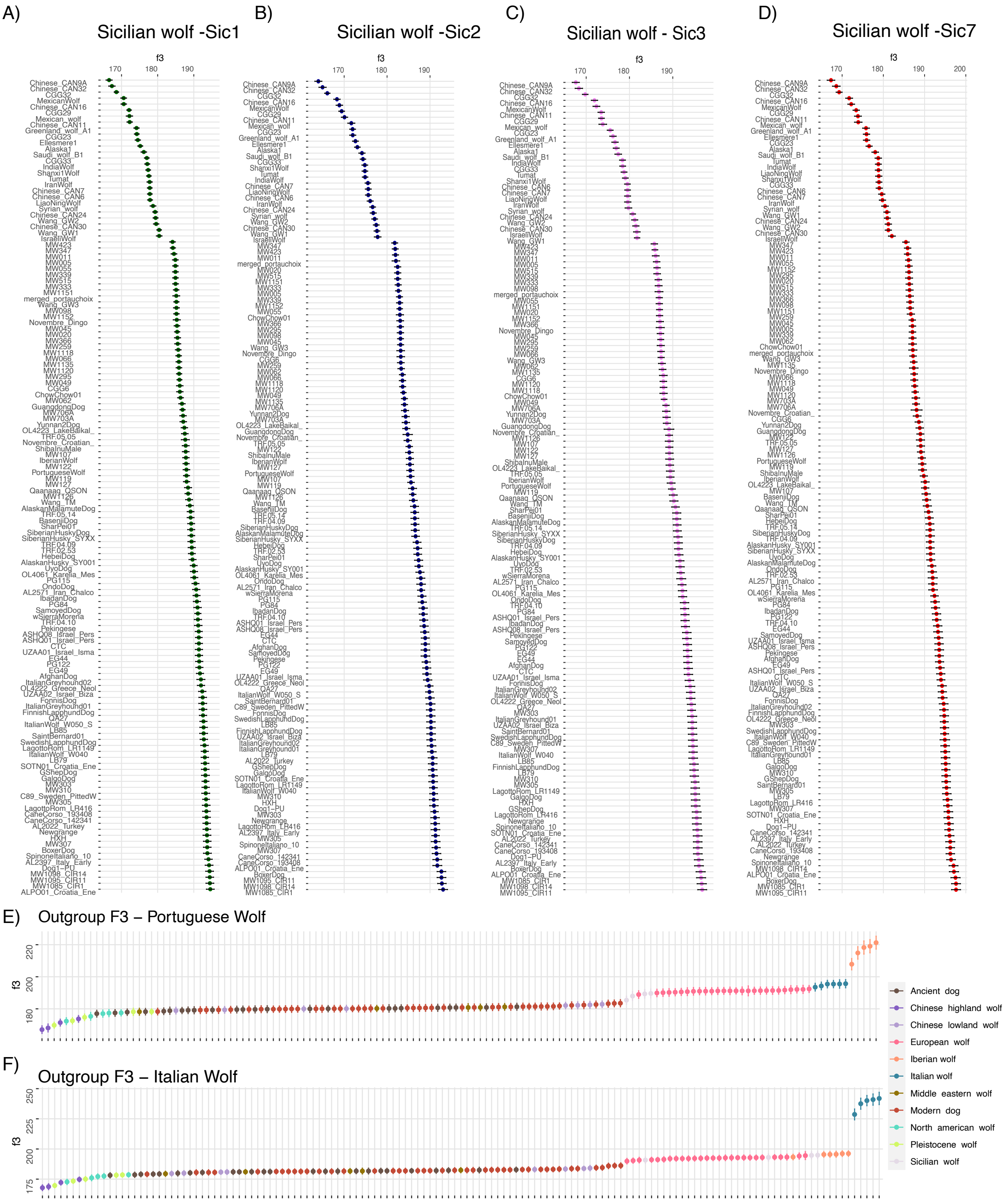


**Figure S3: Outgroup f3 for Sicilian wolves and Portuguese wolf**

A-D) Outgroup f3 of the four Sicilian wolves (Sic1, Sic2, Sic3 and Sic7).

E-F) Outgroup f3 of the Portuguese wolf and Italian wolf as comparison.

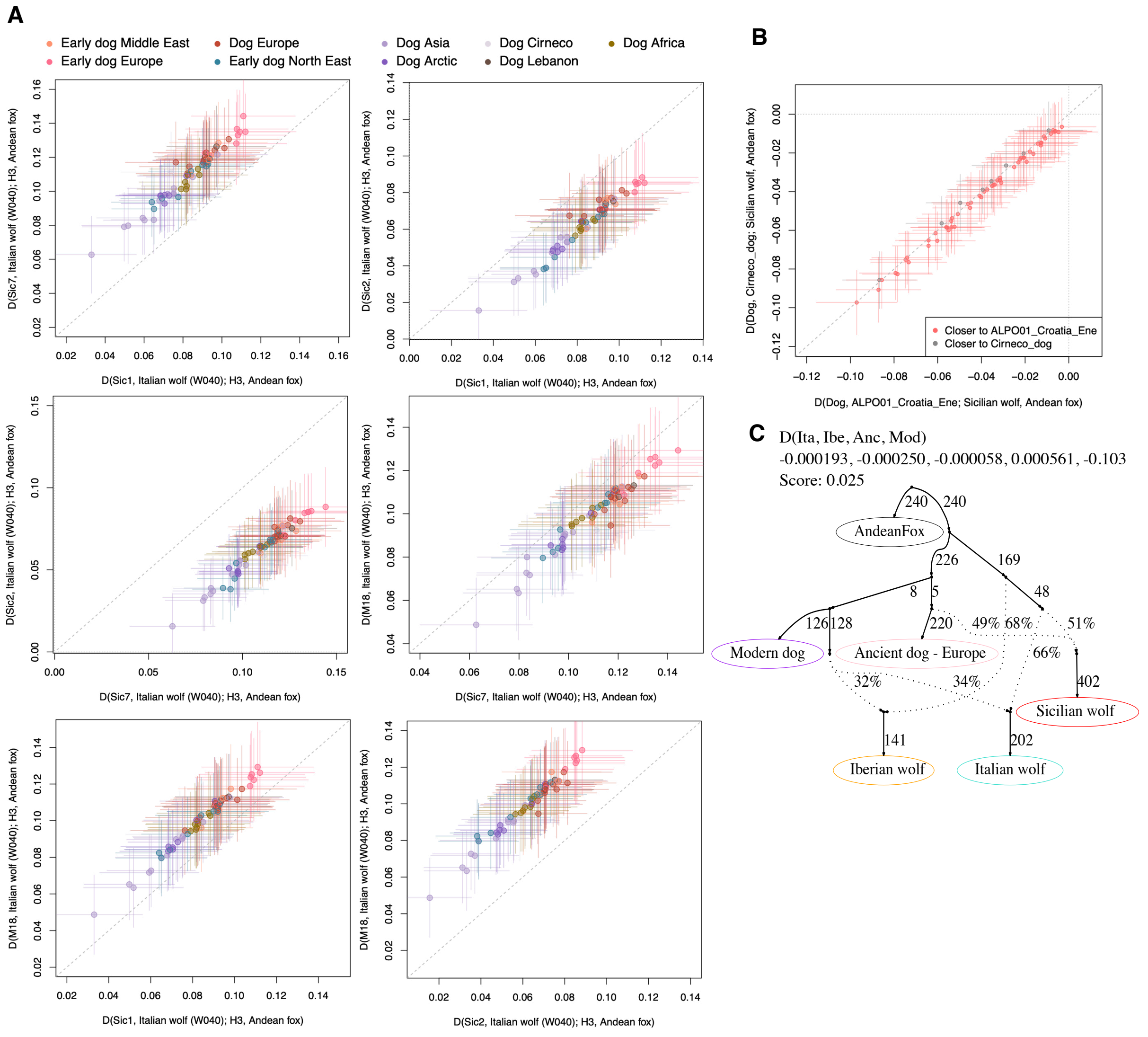


**Figure S4: D-statistic testing for gene flow between Sicilian wolves and dogs using Admixtools [7].**

A) Combination of D-statistics in the form of D(Sicilian wolf, Italian wolf, *Dogs*, Andean fox) both on the x- and y-axes, computed using qpDstats of Admixtools. All tests resulted in significant Z scores (| Z | ≥ 3.3). All the Sicilian wolves result to be closer to ancient dogs (in pink) while Sic3(M1) and Sic7 show to retain the highest dog component.

B) D-statistics test of the form D(*Dogs*, Croatia Eneolithic dog (ALP01); Sicilian wolf, Andean fox) on the x axis and D(*Dogs*, Cirneco dog; Sicilian wolf, Andean fox) on the y axis, computed using qpDstats of Admixtools. All tests resulted in significant Z scores (| Z | ≥ 3.3). The results confirm allele sharing between Sic1 and European wolves, showing a higher signal (D>0 and Z>3) when Italian wolves are in H3.

C) qpGraph admixture modeling the Sicilian wolf as a mixture between the common ancestor of the Italian wolf and ancient European dogs.

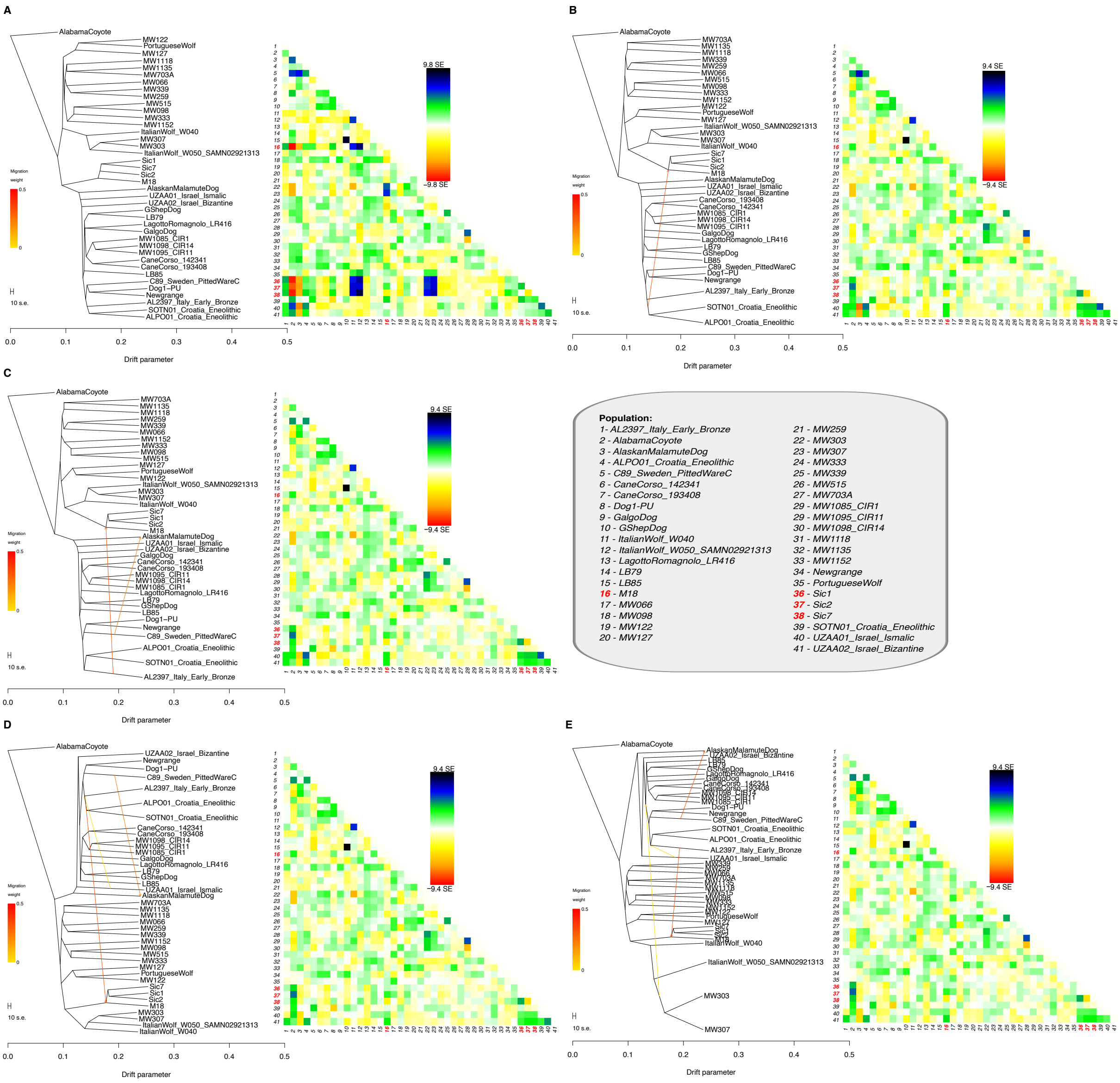


**Figure S5: TreeMix [8] results including 0 up to 4 migration events.**

A-E) TreeMix outputs with 0 to 4 migration edges and their residuals. A) TreeMix tree without migration edges shows the Sicilian wolves basal to modern and ancient dogs. B) TreeMix tree with one migration edge shifts the Sicilian wolves position as sister to the Italian wolves and a first arrow (with high weight) departs from the two Croatian Eneolithic dogs into the base of the Sicilian wolves. C) TreeMix tree with two migration edges: a second arrow from the Swedish Mesolithic dog (C89) goes into the Alaskan Malamute. D) TreeMix tree with three migration edges: the third migration edge brings a Middle eastern component from a ~900BP Israelian dog into Southern European ancient dogs (Italy and Croatia). E) TreeMix tree showing four migration edges: the fourth arrow goes from two Italian wolf samples (MW303 and MW307) to the base of modern dogs.
